# A Piezo1 Open State Reveals a Multi-fenestrated Ion Permeation Pathway

**DOI:** 10.1101/2020.03.12.988378

**Authors:** Wenjuan Jiang, John Smith Del Rosario, Wesley Botello-Smith, Siyuan Zhao, Yi-chun Lin, Han Zhang, Jérôme Lacroix, Tibor Rohacs, Yun Lyna Luo

## Abstract

Force-sensing Piezo channels are essential to many aspects of vertebrate physiology. Activation of Piezo1 is facilitated by the presence of negative membrane lipids in the inner leaflet, such as phosphatidylinositol-4,5-bisphosphate (PIP_2_). Here, to study how Piezo1 opens, we performed molecular dynamics simulations of Piezo1 in membranes flattened by the periodic boundary effect and with or without PIP_2_ lipids. The Piezo1 pore spontaneously opens in the asymmetrical bilayer but not in the symmetric membrane or when PIP_2_ lipids are neutralized. Electrophysiological characterization of putative PIP_2_-interacting Piezo1 residues suggests the contribution of multiple PIP_2_ binding sites. Our Piezo1 open state recapitulates ionic selectivity, unitary conductance and mutant phenotypes obtained from numerous experimental studies. Tracking ion diffusion through the open pore reveals the presence of intracellular and extracellular fenestrations, delineating a multi-fenestrated permeation pathway. This open state sheds light on the mechanisms of lipid modulation, permeation, and selectivity in a Piezo channel.

## INTRODUCTION

Piezos are homotrimeric mechanosensitive channels expressed at the plasma membrane of many cell types in vertebrate animals. They transduce various forms of mechanical stimuli, such as fluid flow or membrane stretch, into electrochemical signals that contribute to a large array of biological functions, including somatovisceral sensation, proprioception, vascular development, blood pressure regulation, osmotic homeostasis, and epithelial growth (*1*). Patients carrying gain or loss of function Piezo mutations present various disease conditions such as xerocytosis, arthrogryposis, loss of proprioception and lymphedema (*2*). Upregulation of Piezo2 activity correlates with inflammation-induced pain states (*3*), whereas gain-of-function Piezo1 variants confer Malaria resistance in humans and in animal models (*4*). The association between Piezo functions and disease states suggests these channels could constitute therapeutic targets for future clinical interventions.

Piezo channels sense mechanical cue transmitted directly from the membrane, and thus obey the so-called force-from-lipid paradigm. High-resolution cryo-electron (cryo-EM) microscopy structures of Piezo1 and Piezo2 revealed a unique molecular architecture consisting of three long peripheral transmembrane domains or arms, and a central region harboring a unique transmembrane pore and an extracellular cap domain (*5-8*). In these structures, the central pore, formed by three inner pore helices, is occluded by hydrophobic side chains and is too narrow to support ion conduction, indicating Piezo1 is captured in a non-conducting state. In this non-conducting conformation, the arms are arranged in tri-dimensional spirals, giving Piezos a triskelion, or propeller-like, shape when viewed perpendicularly to the membrane plane and a bowl-like shape when viewed parallel to it. This curvature around the Piezo arms creates a local curvature, or dome, in the lipid bilayer, suggests the arms sense mechanical forces transmitted from lipids by sensing tension-induced flattening of the membrane (*6, 9, 10*).

Using all-atom (AA) molecular dynamics (MD) simulations, we have recently shown that a truncated Piezo1 computational model spontaneously creates the lipid dome in a relaxed (zero tension) POPC membrane (*11*). We also showed that the dome rapidly flattens when membrane tension is gradually increased. Despite flattening of the arms, however, the pore did not open. Since the Piezo1 arms are anticipated to act as mechanical levers, the shorter arms in our truncated model may reduce the output force (on the pore) to the input effort (arm motion). Another possibility for the absence of opening motions in the pore may have come from the symmetric property of our simulated bilayer: indeed, electrophysiological recordings showed Piezo1 remains fully closed when reconstituted in symmetrical bilayers but spontaneously opens in asymmetric bilayers containing dioleoyl-sn-glycero-3-phosphatidic acid (DOPA) or lysophosphatidic acid (LPA) in the inner leaflet (*12, 13*). DOPA and LPA differ in the number of fatty acid tails but are both negatively-charged. Interestingly, the presence of negatively-charged PIP_2_ or phosphatidylserine (PS) lipids in the inner leaflet also promote channel activation (*14-16*). We thus reasoned that adding the missing arm regions and adding negatively charged PIP_2_ lipids in the inner leaflet will allow us to computationally capture a Piezo1 open state.

## RESULTS

### Piezo1 clustering induces flattening of the Piezo arms

Our previous proof-of-concept AA simulation showed that the membrane curvature, or lipid dome, imposed by the resting conformation of a truncated Piezo1 takes place over 3 μs (*11*). To reduce computational time, here we first used a 12 μs Coarse-Grained (CG) Martini simulation to enable rapid lipid diffusion and dome formation while the Piezo1 backbone was kept rigid. We performed these CG simulations on both a symmetrical POPC membrane (PC:PC) and an asymmetrical POPC membrane containing 5% PIP_2_ in the inner leaflet (PC:PC/PIP_2_) **(Figure 1a**, systems I and II). The CG systems were then mapped back to an AA system and simulated it for an additional 2 μs (**Figure 1a**, systems III and IV). In all MD simulations with explicit solvent, the periodic boundary conditions (PBC) create an infinite lattice where the simulated system is infinitely replicated throughout virtual space. The PBC thus creates a virtual cluster of channels where proteins occupy about 32% of the total membrane area (see **Table S1** for system details).

**Figure 1.**
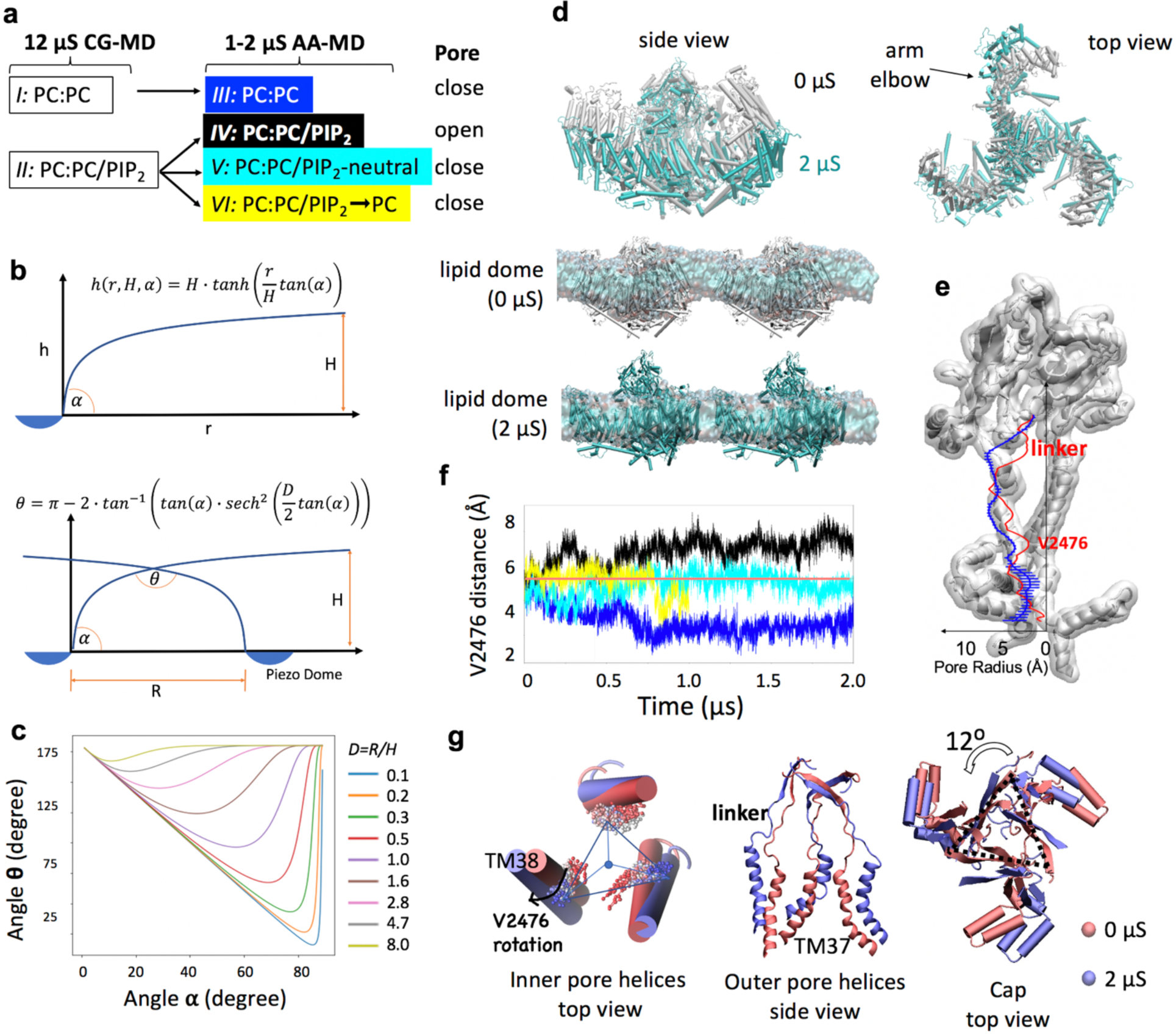
Spontaneous opening of Piezo1. (**a**) Summary of simulated CG systems and AA systems (**Table S1** for details). (**b**) Hyperbolic tangent model, in which *H* is the distance from Piezo dome to the bulk membrane midplane; *R* is the distance between two Piezo dome perimeters; *D* is the separation ratio *R/H*; ***α*** is the angle of Piezo1 arm inclination; ***θ*** is the angle of membrane footprint intersection (see Methods for details). (**c**) The biphasic relationship between ***α*** and ***θ***, and the dependence on the Piezo separation ratio *D*. (**d**) Overlap of a single Piezo1 backbone at 0 μs (white) and 2 μs (cyan) of AA simulation of PC:PC/PIP_2_ system; flattening of the lipid dome between two Piezos, illustrated by the snapshots at 0 and 2 μs of AA simulation of PC:PC/PIP_2_ system (see **Figure S1** for the time evolution of arm flattening in both PC:PC and PC:PC/PIP_2_ systems). (**e**) Pore radius profile calculated using initial atomic coordinates (red) and the last five atomic coordinates taken from the 2 μs AA simulation (1 ns apart) of the PC:PC/PIP_2_ system (blue). Error bars are standard deviation. Radius profiles are overlaid on the Piezo1 pore surface, with one of the three subunits removed for clarity. The position of V2476 and linker region are indicated on the radius profile. (**f**) The time evolution of the radius of the Piezo1 hydrophobic gate, calculated as the nearest vertex to the centroid of three V2476 residues. The color code of the four systems is shown in panel a. The red straight line indicates the distance measured from original cryo-EM structure. (**g**) Comparing the conformations of Piezo1 inner pore helices (three TM38), outer pore helices (three TM37), and cap domain at 0 μs (red) and 2 μs (blue) of AA simulation of PC:PC/PIP_2_ system. The rotation of the hydrophobic gate residue V2476 is illustrated by overlapping the V2476 sidechain trajectories from red to blue. The linker region that connects the cap beta-sheet (residue G2193-G2234) with TM37 is also shown.

The membrane deformation induced by a single Piezo1 channel (membrane footprint) extends well beyond the local dome and decays within tens of nanometers (*9*). Hence, when neighboring channels are closer than this distance, their footprints overlap with an angle smaller than 180 degrees, thus creating an additional energy penalty for membrane deformation. To study how this footprint overlapping affects the lipid dome and the conformation of the arms, we used a simple hyperbolic tangent model (**Figure 1b**, see Methods for details). This model suggests a biphasic behavior between footprint overlap angle (***θ***) and the inclination of the Piezo1 arm (dome angle ***α***) (**Figure 1c**). When the dome angle ***α*** is smaller than a critical value, the footprint flattening (increasing ***θ***) leads to a flattening of the arms (decreased ***α***). In contrast, when the dome angle is larger than this critical angle, the overlap flattening leads to curving the arms even more (increased ***α***).

In our system, the closest distance between neighboring Piezo1 arms (***R***) is around 3 nm. The height from the dome apex to the bulk membrane in absence neighboring channels (***H***) is predicted to be around 14 nm (*9*). Thus the separation distance ***D=R/H*** is less than 1 in our simulated systems. Using these parameters, the hyperbolic tangent model predicts that the critical value of the dome angle ***α*** is above 60 degrees (**Figure 1c**), which is larger than the 30° dome angle determined from cryo-EM structures. As expected for a dome angle lower than the critical value, we observed a spontaneous flattening of the overlap footprint of the lipid dome and of the Piezo1 arms in both PC:PC and PC:PC/PIP_2_ bilayer systems (**Figure 1d and Figure S1**). During flattening, the arms also undergo a counterclockwise twist, mimicking a blooming-like motion (**Figure 1d** top view).

### Structural changes associated with pore opening

To track the pore opening, we monitored the size of the narrowest region of the pore along our AA trajectories. This region corresponds to the position of valine 2476, which has been proposed to form a hydrophobic barrier (*17*), occluding the pore in cryo-EM structures (**Figure 1e**). In the symmetric PC:PC membrane, the radius of the hydrophobic barrier decreases during the 2 us simulation from about 5 Å to 2 Å, constricting the pore even further than in cryo-EM structures (**Figure 1f**, system III: PC:PC, blue trace). In contrast, in the asymmetrical membrane, the radius of the valine barrier increases from 5 Å to 7 Å during the first 750 ns and remains above 7 Å for the remainder of the simulation **(Figure 1f**, system IV: PC:PC/PIP_2_ in black). The widening of the hydrophobic barrier correlates with an outward tilt of the intracellular end of the inner pore helices (TM38). This tilt rotates the valine 2476 side chains away from the pore lumen, increasing its diameter (**Figure 1g**). The outer pore helices (TM37) have a larger degree of outward motion (**Figure 1g**). As a result, the pore radius at the linker region between the cap and TM37 increased (**Figure 1e**). In addition, the cap domain shows on average 12 degrees of counter-clockwise rotation.

### PIP_2_-mediated electrostatic interactions favor the open state

To determine the contribution of the negative charges of PIP_2_ to pore opening, we performed a second control simulation where all these charges are computationally silenced (**Figure 1a**, system V: PC:PC/PIP_2_-neutral). In this new system, the Piezo1 pore opening was not observed as the radius of V2476 remains the same as in cryo-EM structure over 2 μs AA simulation (**Figure 1f**, cyan trace). A reduction of the overall binding interactions between charge-neutralized PIP_2_ lipids and the Piezo1 channel further confirms the electrostatic nature of these interactions (**Figure S2**). To rule out the contribution of non-electrostatic differences between PC:PC and PC:PC/PIP_2_ simulations (such as different amino acid side chain orientations introduced by distinct CG simulations or different lipid number), we conducted a third control simulation where all 39 PIP_2_ lipids from system IV are replaced by the same number of POPC molecules while the rest of the system is kept identical. In this system (**Figure 1af**, system VI: PC:PC/PIP_2_→PC, yellow trace), the pore remained closed. Taken together, the fact that all three control systems (III, V, and VI) failed to open the pore strongly suggests that PIP_2_-mediated lipid-protein electrostatic interactions facilitate Piezo1 opening.

### Validations of our computational Piezo1 open state

In cryo-EM Piezo1 structures, hydrophobic cavities are clearly seen above and below the narrow valine pore constriction. Through these conduits, POPC tails penetrate into the pore lumen during our backbone-restrained CG simulations (**Figure S3)**. Such pore occlusion by membrane lipids is not uncommon and has been proposed to participate in a bona fide physiological gating in mechanosensitive MscS and TRAAK channels (*18-21*). Since the time needed for these lipids to spontaneously leave the pore may extend beyond our 2 μs simulation, all pore lipids were deleted at the end of the trajectory, allowing water and ions to diffuse through the pore (**Figure 2a**). This permits us to calculate the unitary ionic conductance through the pore.

**Figure 2.**
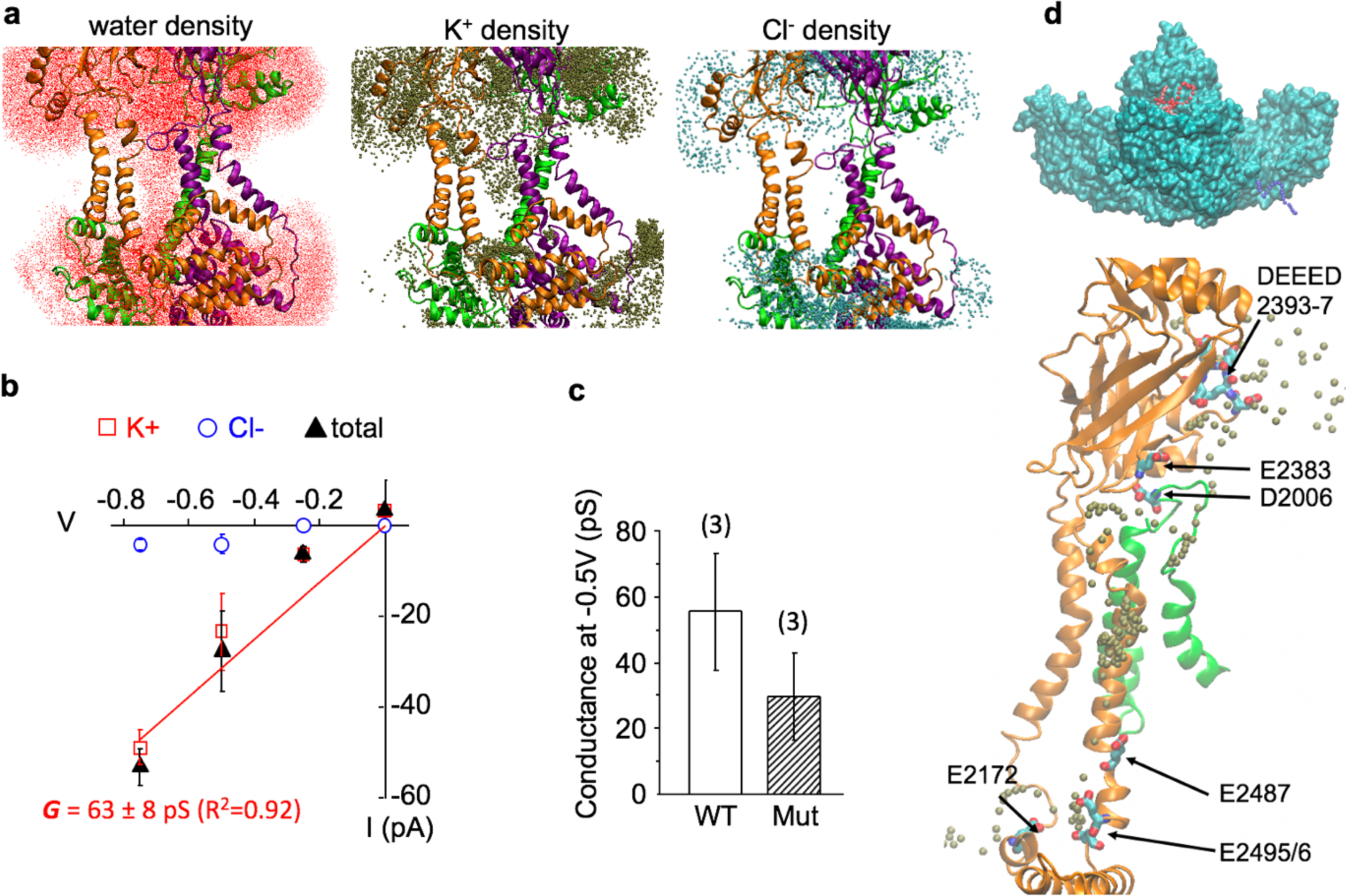
Calculated conductance and multi-fenestrated permeation pathway of Piezo1 channel. (**a**). Water and ion density in the Piezo1 pore from 150 ns PC:PC/PIP_2_ simulation at −500 mV voltage. The protein backbone is colored by subunits (orange, purple, and green). Only inner and outer helices, part of cap and CTD domains are shown for clarity. (**b**). The total ionic conductance (black triangles) and individual conductance of K^+^ (red open rectangles) and Cl^-^ (blue open circles) ions obtained from AA-MD simulation of WT Piezo1. The red line is a linear fit of the total current. (**c**). Unitary conductance of WT Piezo1 (WT) and of Piezo1 where the negative charge of E2133 is neutralized (Mut) during three independent 50 ns simulations at V = −500 mV. (**d**). Representative multi-fenestrated permeation pathway (see **Video1** for single event, and **Video2** for cumulative density). Top: a 9.5 ns trajectory of a single K^+^ ion colored in timestep from red to blue, with the whole PIEOZ1 shown in cyan surface. Bottom: High K^+^ density hotspots are shown on the protein backbone of a single subunit pore region (orange), except D2006 which is located in the loop of a nearby subunit (green). The K^+^ is colored in brown. Hotspot residues are shown in licorice with atom color code (red oxygen, cyan carbon, blue nitrogen).

The unitary conductance of ion channels is experimentally obtained by fitting the slope of the current vs. voltage relationship obtained from single-channel recordings. To calculate the conductance from MD simulation, constant electric fields corresponding to the transmembrane potentials of −250, −500, and −4750 mV were applied perpendicular to the membrane to all the atoms in the simulation box, in the presence of symmetrical 150 mM KCl concentration. For each voltage, three consecutive 50 ns simulations were carried out to calculate the mean and standard error of the K^+^ and Cl^-^ ions permeation events. A membrane tension of 14.2 mN m-1 (about −10 bar considering water box fluctuations) was found to be optimal to stabilize the open pore conformation throughout the trajectory at all voltages (**Figure S4** pore RMSD). The total ionic current was determined by calculating the displacement of all charges across the membrane. A least-square fitting of the I-V curve, subjecting to the constraint of zero reversal potential at symmetric salt concentration, yields the conductance of 63±8 pS (n=3) (**Figure 2b**), in excellent agreement with the experimentally-obtained conductance of 60 pS in the absence of divalent cations (*22*).

In addition, the Piezo1 pore remains cation-selective across all tested voltages (**Figure 2b**). Using the non-zero Cl^-^ permeation events at −500 and −750 mV simulations, we obtained a K^+^:Cl^-^ permeation ratio in the range of 1:5 to 1:13 (n=3). These approximated values are remarkably similar to the reported Na^+^:Cl^-^ permeation ratios of 1:7 and 1:13 for mouse Piezo1 (*23, 24*).

We further tested whether our open state can reproduce the phenotype of a conductance-reducing mutant. The conserved glutamate 2133 residue located in the anchor region is an important determinant of channel conductance as charge neutralization mutations E2133A and E2133Q produce a two-fold reduction of unitary conductance (*23*). Using the open state conformation, we computationally silenced the negative charge of E2133. As expected, this charge neutralization reduced the frequency of permeation events for both K^+^ and Cl^-^ ions, leading to a nearly two-fold reduction of the unitary conductance from 55±8 pS to 30±6 pS (n=3) (**Figure 2c**).

### Multi-fenestrated ion permeation pathway and cation-selective residues

In our previous AA simulation of a truncated Piezo1, we noticed the presence of K+ ions within three intracellular fenestrations and in an intracellular pore vestibule located underneath the position of V2476. Here, the K^+^ permeation pathway captured under the electric field not only confirmed these intracellular fenestrations but also revealed that the extracellular ions enter the pore via wide lateral fenestrations located between the extracellular mouth of the pore and the pore-facing interface of the cap (**Figure 2d, Video 1 and 2**). The K^+^ density from conductance simulations revealed several hotspots along the ion permeation pathway, indicating longer K+ residence time. The rim surrounding the entryway for extracellular fenestrations contains the negatively-charged residues DEED 2393-7 (DEED loop), E2383 in the cap and D2006 located in the arm in close proximity to the cap. Several negatively-charged residues are located along the narrower intracellular entryway, such as E2172 on the anchor, and E2487, E2495/6 on the inner helix (TM38) (**Figure 2d**). Experimental neutralization of many of these residues (2393-7, E2487, E2495/6) significantly reduced or abolished cation selectivity strongly supporting the twisted ion permeation pathway unraveled by our simulations (*23, 24*).

### The Piezo1 open state is consistent with intersubunit distance constraints

A recent study showed that inserting an intersubunit cysteine bridge between cap residues A2328 and P2382 prevents the opening of Piezo1 by cell indentation (*25*). The same phenotype was observed when a disulfide bridge is inserted between the cap residue E2257 and the arm residue R1762. These experimental results showed that for both pairs of residues, the inter-residue distance permits disulfide bond formation in the close state but not in the open state. We hence sought to confirm whether those two inter-residue distances are within disulfide bond formation at the beginning of our PC:PC/PIP_2_ simulation and increase beyond disulfide bond formation during the course of the simulation. As expected, the three intersubunit A2328-P2382 distances increased from 5 Å to 8 ∼ 18 Å (**Figure 3**). The A2328-P2382 pairs are located at the base of the cap, which is linked with the outer pore helices (TM37) through a linker region. The simulation trajectory shows that when the arms flatten, the base of the cap widens to enable outward motion TM37 helices (**Figure 1eg**). This widening cap motion separates A2328 and P2382 beyond disulfide bond formation. In addition, as expected during the flattening of the arms, all three E2257-R1762 distances between cap and arm increased to more than 20 Å during the trajectory (**Figure 3**). In the closed state, the cap motion is prohibited by the close contact with the arms. Thus, the cap rotation shown in **Figure 1g** is only allowed by arm flattening which breaks the cap and arm contact. Both intersubunit distances are located in the bottom part of the cap, which is consistent with the fenestration observed below the cap. The K^+^ pathway suggests that the extracellular cations are guided into the upper vestibule of the pore by the DEED loop (residues 2393-7) on the surface of the cap that reaches out to the bulk region, and then pulled down by E2383 on the bottom of the cap domain and D2006 on the loop of Piezo arm under the cap (**Figure 2d**).

**Figure 3.**
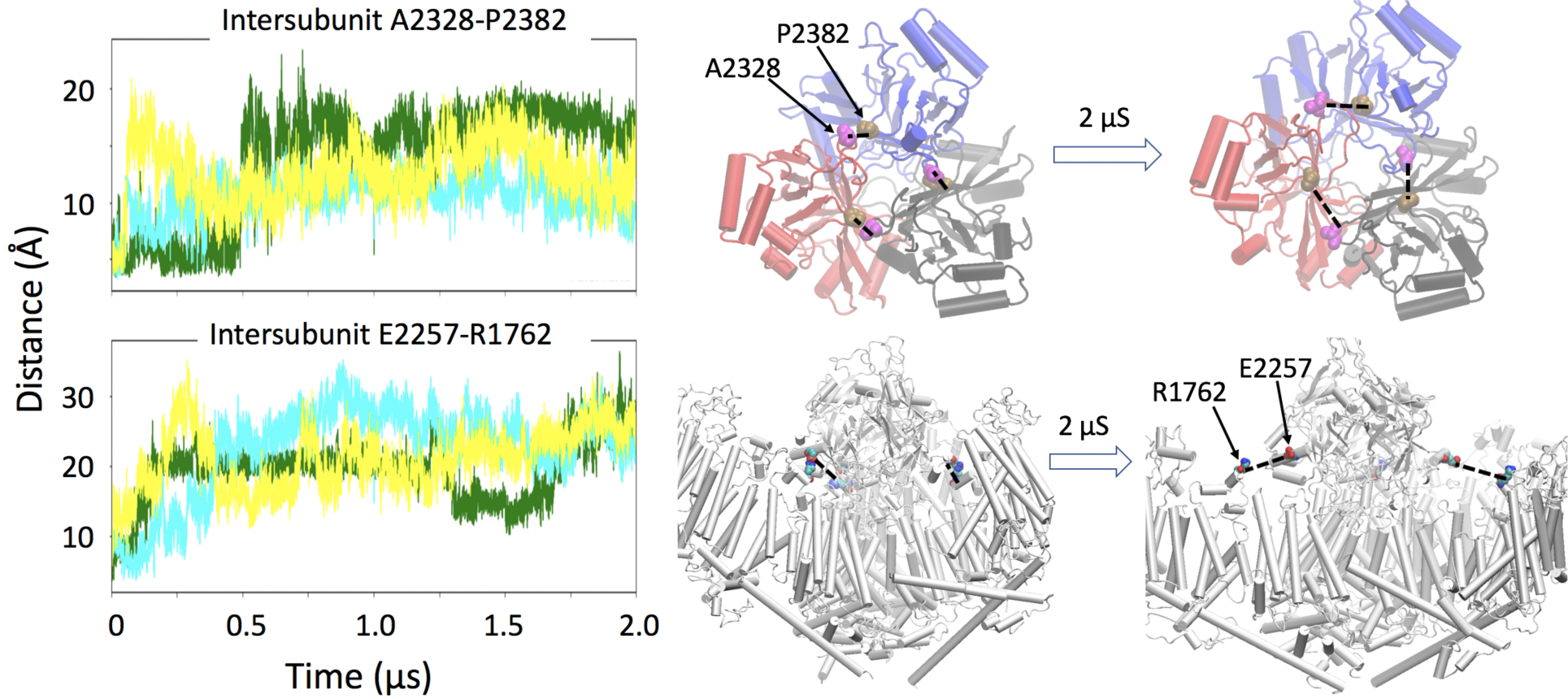
Conformational changes during Piezo1 activation are consistent with inhibitory disulfide bridges. **(a).** Distance between β carbon of cross-linking residues during the simulation time. Three colors represent three pairs of intersubunit residues. **(b).** The increased distances illustrated on the protein structure at the beginning and end of the AA simulation.

### Mutations of putative PIP_2_ binding sites recapitulate PIP_2_ depletion phenotype

A large number of PIP_2_ binding and unbinding events from microseconds of CG simulation and AA simulation of Piezo1 in PC:PC/PIP_2_ bilayer allow us to identify the PIP_2_ binding hotspot on Piezo1. Out of 143 cationic residues in each subunit of the current Piezo1 model, only 16 residues show at least one PIP2 bound over 60% of 12 μs CG simulation and over 90% of 2 μs AA simulation time (844, 948/9, 1023-6, 1727/8, 2040/1, 2113, 2182-5 in **Figure 4a)**. The binding distance is determined by the first minimum from the radial distribution function between arginine/lysine sidechains and the PIP_2_ headgroups (see Methods). Those 16 residues are located at three different regions of the intracellular protein surface, namely the peripheral arm region, the convex side of the arm region, and the pore and anchor region (**Figure 4b)**.

**Figure 4.**
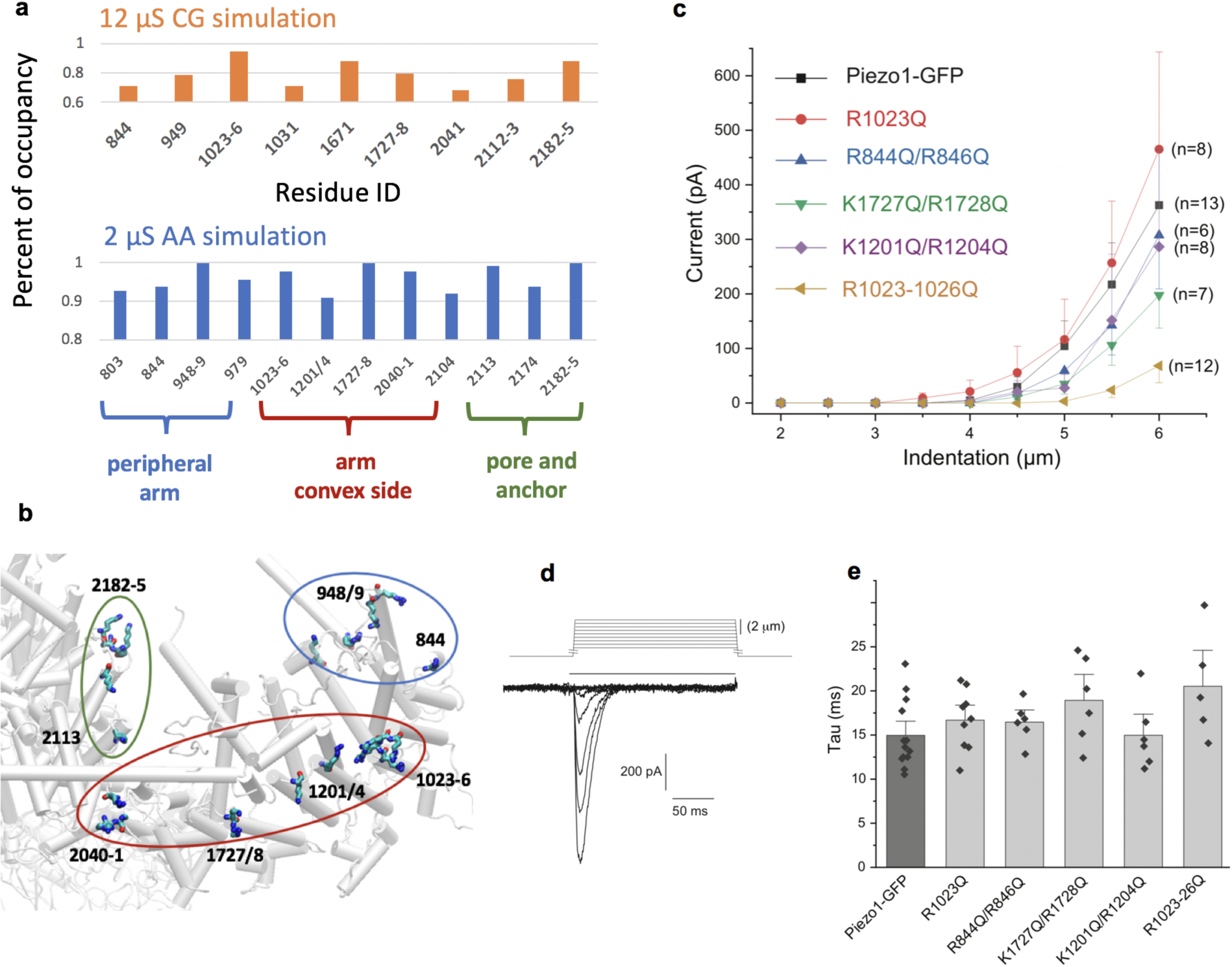
The effect of mutating PIP_2_ interacting residue clusters on Piezo1 channel activity. (**a).** The residues in mouse Piezo1 that have maximum PIP_2_ occupancy above 60% of 12 μs CG simulation or above 90% of 2 μs CG simulation. (**b).** Hotspot residues clustered by locations on the Piezo1 structure (bottom view). (**c).** Summary of current amplitudes in response to increasing mechanical stimuli for wild type and mutant Piezo1 channels. Whole cell patch clamp experiments on HEK293 cells transfected with the GFP-tagged mouse Piezo1 and its mutants were performed as described in the methods section. (**d).** Representative trace of wild type Piezo1 currents in response to increasing mechanical indentations. **(e).** Summary of the inactivation time constant for wild type and mutant Piezo1 currents.

To investigate whether some of those PIP_2_ bindings are functionally relevant, we generated R/K to Q mutations to neutralize the positive charge and thus abolish PIP_2_ binding. Nearby cationic residues were mutated together to ensure a complete loss of PIP_2_ binding at that particular position. At the peripheral arm region (**Figure 4b**), we tested a double mutant R884/6Q, which showed a minimal reduction in current amplitudes (**Figure 4c)**, and excluded the residues on the concave side of the peripheral arm as the missing sequence on the N-terminal region may lead to an overestimation of the PIP_2_ binding. On the convex side of Piezo1 arm, PIP_2_ binding hotspots spread the whole arm region. The double mutant K1201Q/R1204Q showed a minimal decrease in MA current amplitudes, while the K1727Q/R1728Q showed a 44% decrease at the maximal stimulation strength (**Figure 4cd**). A single mutant R1023Q at the elbow region of the Piezo1 arms showed MA currents similar to that in wild type channels, however the quadruple mutant R1023-6Q showed a current reduction by ∼80 %. This construct however showed visibly dimmer GFP fluorescence, therefore some of the decrease was likely due to decreased expression levels. Overall, none of the mutations abolished Piezo1 function, indicating that none of the putative individual binding sites is indispensable for mechanical activation of Piezo1. It is also consistent with previous results showing that depletion of PIP_2_ in a cellular context does not completely abolish Piezo1 activity (*14*). None of the mutations changed the inactivation time constant significantly (**Figure 4e**).

## DISCUSSION

The open state generated from our MD simulation not only faithfully reproduces unitary channel conductance but also recapitulates the selectivity of monovalent cations vs. anions (**Figure 2b**). We show that the main activation gate is located near V2476, as anticipated from non-conducting Piezo1 structures. Pore opening is associated with an outward motion of outer pore helices, allowing tilting of inner helices and rotation of V2476 side chains away from the pore lumen (**Figure 1g**). In addition, we observed a concomitant rotation and widening of pore-facing cap sub-domains. Interestingly, rearrangements of pore-facing sub-regions of the cap have been recently shown as necessary for Piezo1 activation (*25*), highlighting the importance of cap flexibility in permitting channel activation (**Figure 3**).

The Piezo1 permeation pathway uncovered here reveals that ions enter the open pore via lateral fenestrations (one per subunit) on both the extracellular and intracellular sides (**Figure 2d, Video 1 and 2**). Lateral fenestrations are not uncommon as they have been identified in many families of ion channels and transporters. The multi-fenestrated permeation pathway of Piezo1 is further consistent with the observation that the deletion of a C-terminal beam-to-latch region of Piezo1, which forms a cytosolic plug under the central pore, does not yield constitutively open channels (*26*). We, however, do not exclude the possibility that part of the disordered loops between beam and repeat A, absent from the cryo-EM structures and our model, may gate or partial gate the intracellular fenestration (*27*). Our MD simulation reveals K^+^ ions interact with basic residues (mainly glutamate) known to contribute to ion selectivity both in the extracellular and intracellular fenestrations as well as residues that have not yet been experimentally tested. Together, these observations indicate that the cation selectivity of Piezo1 is governed by multiple residue-ion electrostatic interactions clustered at both intracellular and extracellular fenestrations.

Our simulations underscore the exquisite interplay between local membrane geometry and Piezo1 curvature, a property highlighted from cryo-electron microscopy and atomic force microscopy studies (*9, 10*). The clustering effect produced by the periodic boundary condition (PBC) of MD simulations mimics a high-density channel cluster and imposes a flattening of the Piezo1-induced membrane footprints, reducing the curvature of the lipid dome and of the Piezo1 arms. The Piezo1 conformational changes induced by channel clustering has two major components: a flattening of the whole arm when viewed parallel to the membrane and a straightening of the proximal N-terminal region (rotation around the elbow) when viewed perpendicular to the membrane (**Figure 1d**). These conformational changes observed here due to membrane flattening are similar to the ones observed from a truncated Piezo1 simulation when applying membrane tension (*11*). There are three helical bundles (Piezo repeats) on the N-terminal peripheral region of the arms not present in Piezo1 cryo-EM structures. It is possible that the longer arms may increase the sensitivity to the membrane tension.

Interestingly, Piezo1 channels seem to form clusters when heterologously expressed in mammalian cells and this clustering has been proposed to play a role in concerted gating transitions such as collective loss of inactivation (*28-30*). While it is currently unclear whether endogenous Piezo1 form native clusters in vivo, these results together suggest that the gating properties of Piezo channels can be modulated by the local channel density at the plasma membrane. According to our hyperbolic tangent model, when the distance between neighboring channels is small, clustering favors flattening of the lipid dome and, consequently, of the Piezo1 arms. On the contrary, if the distance between two neighboring channels becomes sufficiently large, clustering is predicted to increase the curvature of the arms (**Figure 1b**). The critical inter-channel distance separating these two scenarios depends on precise geometric parameters of the Piezo1-induced membrane footprint, which also depends on bilayer rigidity and membrane tension (*9*).

Crowding-induced membrane footprint flattening may not be the only possible mechanism underlying concerted gating in clustered Piezo channels. Cooperative gating may also be governed by direct protein-protein interactions between nearby channels or by indirect interactions mediated by auxiliary proteins (*31, 32*). Changes in bilayer thickness due to hydrophobic mismatch has been proposed to induce cooperative gating between neighboring MscL mechanosensitive channels, and thus could contribute to cooperative gating in Piezo channels. Other entropic contributions due to reduced membrane fluctuations in Piezo clusters may also collectively influence gating property. Whether and how those factors contribute together to the cooperative gating in Piezo clusters with different densities is of interest for further studies. Future MD simulations using PBC aimed at studying Piezo1 clustering may lead to several caveats. First, due to the large size of molecular systems simulating Piezo channels and their large membrane footprints, it will be technically challenging to generate a microsecond-long trajectory of a low-density Piezo1 cluster using available computing resources. Second, channels replicated under PBC condition are mirror images of each other. Thus, MD simulations under PBC cannot replicate spatial heterogeneity of channels in a real cluster. The predicted biphasic behavior of Piezo channels under different cluster densities may be better investigated using high-resolution biophysical approaches, such as high-speed atomic force microscopy or electron cryo-electron microscopy.

Our MD simulations show that the presence of negatively-charged PIP_2_s is necessary for spontaneous Piezo1 pore opening in a 2 microseconds temporal window. This observation is consistent with the spontaneous opening of Piezo1 observed in droplet bilayers containing negatively-charged lipids in the inner leaflet (*12, 13*) and with the increase of the mechanical threshold for Piezo1 activation (reduction of open probability) observed in PIP_2_-depleted cell membranes (*14*). In symmetric PC:PC bilayers, mechanical stress alone is sufficient to activate Piezo1, which indicates that PIP_2_ is not an absolute requirement for Piezo1 activation (*13*). Together, these results suggest PIP_2_s facilitate channel activation, likely by reducing the free energy difference between closed and open states. The Gibbs free energy change associated with the opening transition can be obtained from the following formula:

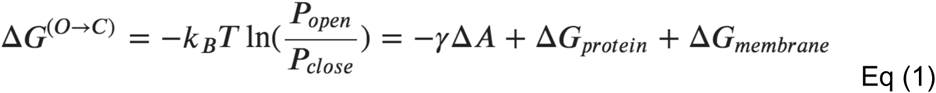

in which *k*_*B*_ the Boltzmann constant, *T* the temperature, *P*_*open*_ and *P*_*closed*_, respectively the probability of the channel being open and closed, *γ* the membrane tension, ΔA the relative change in the membrane surface footprint associated with channel opening, ΔG_protein_ the free energy of protein conformational change in absence of membrane tension, and ΔG_membrane_ the free energy of membrane deformations. Piezo1 clustering could modulate membrane deformability in absence of tension (ΔG_membrane_), therefore altering the open/closed equilibrium.

According to equation (1), PIP_2_s may modulate the closed-open equilibrium of a mechanosensitive channel in several ways. First, they may alter the mechanical properties of bilayer, such as bilayer bending rigidity and bilayer area compressibility. These mechanical constants quantify the energetic cost associated with membrane deformation ΔG_membrane_. Second, the interactions between PIP_2_s and Piezo1 residues may reduce the protein conformational energy ΔG_protein_ by destabilizing the closed state and/or stabilizing the open state. In our simulations, membrane flattening occurs regardless of the presence of PIP_2_ lipids in the membrane. In addition, in our PC:PC/PIP_2_ simulation, the majority of PIP_2_ lipids remain located within atomic proximity to Piezo1, not in the bulk membrane (**Figure S2**). Hence, while we cannot rule out the possibility that PIP_2_ lipids mediate their effects on Piezo1 by changing membrane mechanical properties, our simulations strongly suggest PIP_2_ lipids promote Piezo1 activation by modulating channel energetics ΔG_protein_ through direct lipid-protein interactions. Abolishing the electrostatic interactions between PIP_2_ and PIEOZ1 through charge neutralization of PIP_2_s resulted in a closed channel in our simulation time.

Our mutagenesis data shows that neutralizing mutations of putative PIP_2_ binding residue clusters had minor effects on mechanically-induced channel activity, and even a quadruple mutant was functional, although it required stronger mechanical stimuli to open. Thus, a large number of lipid-protein interactions seem to be necessary to shift the open probability of a correspondingly large membrane protein. The binding of multiple PIP_2_s along the convex side of each arm may amplify the force transmitted from the lipid bilayer to the protein, reducing the tension value (*γ*) required to open the pore. This hypothesis is supported by the fact that stronger mechanical stimuli are required to open Piezo1 when membranes are PIP_2_-depleted (*14*) or when putative PIP_2_ binding sites are mutated (**Figure 4c**). On the other hand, PIP_2_s may help anchor the intracellular side of the arms to the inner leaflet, allowing a better mechanical coupling upon membrane deformation. Such mechanical coupling may reduce the entropic cost associated with the conformational rearrangement of the arms, an effect explained by the population-shift theory of allostery (*33, 34*). Finally, in addition to the change in the thermodynamic quantity, PIP_2_ may also affect the kinetic aspect by reducing the free energy barrier for Piezo1 activation, enabling spontaneous opening in our microsecond MD simulation. However, there is no indication that Piezo1 activation kinetics slow down upon PIP_2_ depletion (*14*). The interaction of PIP_2_ with multiple binding sites on a large protein surface area of Piezo1 is in contrast to the binding of PIP_2_ to well defined single binding sites per subunit in several other PIP_2_ regulated ion channels, such as inwardly rectifying K^+^ channels (*35*), TRPV5 channels (*36*), and TRPM8 channels (*37*). In TRPV5, the PIP_2_-bound structure shows an open conformation (*36*), compared to PIP_2_-free structures, indicating that binding of a single PIP_2_ molecule per subunit is capable of inducing a conformational change to open the channel. Regulation of Piezo1 by PIP_2_ through multiple binding sites is likely to be far more complex. Mutations of putative PIP_2_ interacting residues had no effect on the inactivation kinetics of mechanically-activated Piezo1 currents (**Figure 4e**). The quadruple Lysine K2182-85 also showed up on our simulations as a cluster frequently interacting with PIP_2_ (**Figure 4a)**. This cluster is equivalent to residues K2166-69 in the human Piezo1 channel. Deletion of these residues is associated with xerocytosis (*38*) and display markedly slower inactivation of mechanically activated Piezo1 currents (*39*). This may indicate that PIP_2_ binding to distinct sites in the channel have different effects on channel function. Since PIP_2_ depletion had no significant effect on Piezo1 inactivation (*14*), it is also possible that the effect of those mutations on channel inactivation is independent of PIP_2_ binding.

In conclusion, the Piezo1 open state generated from MD simulation produces biophysical properties consistent with a large body of published experimental data on unitary single channel conductance, selectivity, and mutant phenotypes. We revealed conformational changes associated with Piezo1 opening, including motions in the mechanosensory domains (arms), outward motions of inner pore helices, twisting motion of Valine residues acting as a hydrophobic gate, and rotation of the extracellular cap domain. These motions are consistent with structure-function studies, including point mutagenesis state-dependent formation of engineered cysteine bridges. The unique multi-fenestrated ion permeation pathway captured from simulations (**Figure 5**) are supported by experimental neutralization of residues along the pathway that reduced or abolished cation selectivity. Reproducing the phenotype of a conductance-reducing mutant provided further validation of this open conformation. In addition, we demonstrate, computationally and experimentally, that Piezo1 opening is facilitated by the presence of electrostatic interactions between PIP_2_ lipids and multiple binding sites located along the Piezo1 arm. With the availability of both Piezo1 open and close states, future computational studies of free-energy landscape of Piezo activation and allosteric network analyses, in combination with experimental studies will be helpful in understanding the molecular underpinnings of lipid modulation on Piezo channels, and provide a new avenue for mechanistic investigation of disease mutations and small molecule drug discovery.

**Figure 5.**
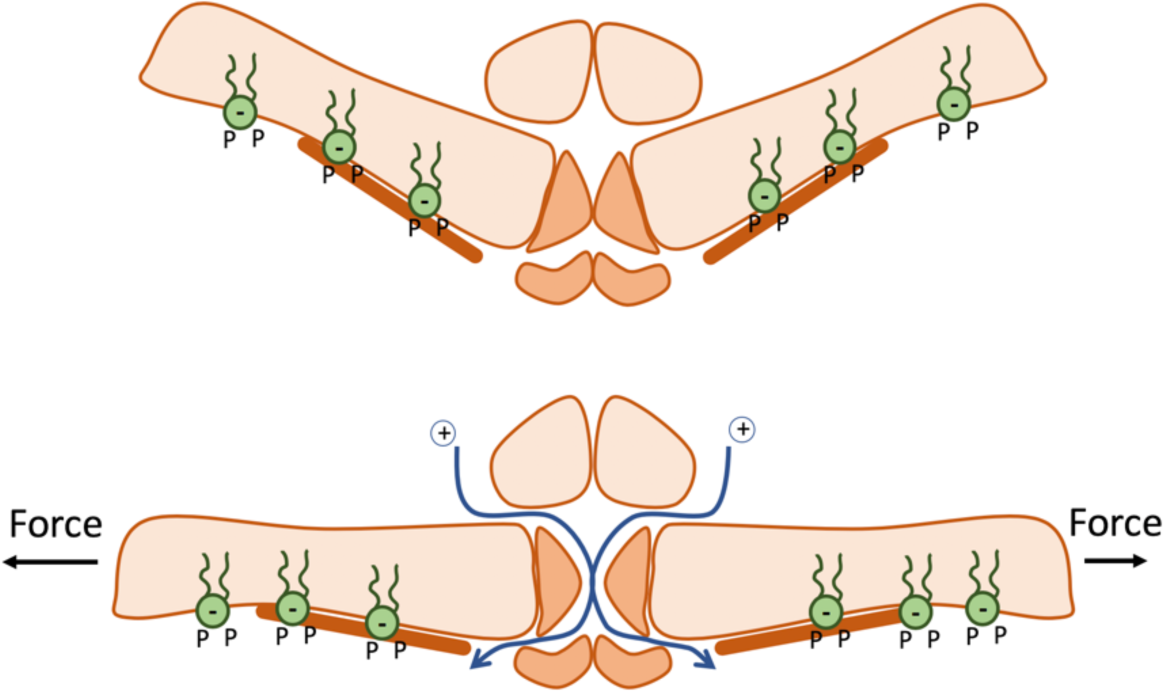
Piezo1 channel activation, facilitated by arm flattening and multiple PIP_2_s binding, reveals a multi-fenestrated ion permeation pathway.

## METHODS

### Derivation of Hyperbolic Tangent Membrane Footprint Model

The hyperbolic tangent model was chosen to mimic two observed properties of the membrane footprint. Firstly, the membrane approaches a flat plateau when distant from the Piezo dome (although ripple-like oscillations/perturbations inherent to such membranes prevent this from being strictly monotonic). This is captured well by the hyperbolic tangent function which monotonically approaches unity as it moves away from the origin (**Figure 1b**). Thus the height of the membrane *H* can serve as a reduced distance unit for the hyperbolic tangent model. Secondly, the membrane footprint exhibits a ‘knee’ like bend (i.e. a point of maximal concavity/curvature). The sharpness at this knee point grows stronger as the angle of inclination of the arms of piezo increases. This can be captured in the hyperbolic tangent model by adding a scaling factor inside the hyperbolic tangent function. More specifically, this scaling constant will be equal to the slope of the hyperbolic tangent model at the origin. Thus, it may be easily related to the angle of inclination of the membrane at the edge of the piezo dome. If the arms of piezo have an angle of inclination *α* with respect to the *xy* plane, the corresponding slope is given as ***tan***^***−1***^(***α***). Lastly, if we place two piezo domes a distance of *D* reduced units apart (note: D=R/H), their membrane footprints will intersect at a minimum distance of *D/2* units apart. Putting this together, we attain the equation (2) below:

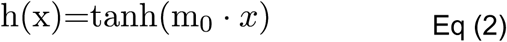

where h(x) is the height of the membrane above the top of the piezo dome at a radial distance of x reduced units away from the edge of the dome, and *m*_0_ is the slope of the membrane at the edge of the piezo dome. Correspondingly, we may calculate the slope of the membrane footprint under this model as in equation (3):

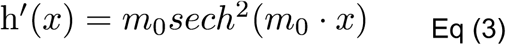

where h’(x) is the slope of the membrane footprint at position x. We may then calculate the slope of the membrane footprint at the point of intersection with another membrane footprint as in equation (4):

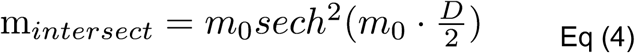

where m_*intersect*_ is the slope of the membrane footprint model at the point where it would intersect another membrane footprint when the edges of the two domes are a distance of *D* reduced units apart. This slope function can be cast as the intersection angle *θ* as a function of the angles of inclination of the membrane footprint at the edge of the piezo dome, *α*. To do so, we first note that if the membranes have a slope of ±*m*_*intersect*_at their intersection then their corresponding angles are the inverse tangent of their slope. This angle is their effective ‘angle of inclination’ at that point, so the angle formed between them would then be pi radians minus their sum (or 180º minus their sum in degrees). This yields equation (5):

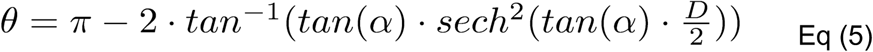

### Coarse-Grained (CG) System Preparation

The CG representation of Piezo 1 was constructed from our previously all-atom mouse Piezo1 model based on the cryo-EM structure (PDB ID 6B3R), which includes residue 782–1365 (Piezo repeat C-F and beam), 1493–1578 (clasp), 1655–1807 (repeat B), and 1952–2546 (repeat A, anchor, TM37, cap, TM38, and CTD) (*11*). Based on these atomistic coordinates, the coarse-grained model using MARTINI v2.2 force field was obtained through the script *martinize.py* and *insane.py* available from the MARTINI web site (*40-42*). For lipids, C16:0/18:1 1-palmitoyl-2-oleoy phosphatidylcholine (POPC) follows the standard Martini 2.0 lipid definitions and building block rule. A modified PI(4,5)P_2_ MARTINI model carrying −4 charge was parameterized to be consistent with the experimental data (see the section below). For all the systems, a 28.4×24.6 nm membrane bilayer was solvated with explicit water in a simulation box of 28.4 × 24.6 × 25.3 nm. 150 mM NaCl was added to each simulation and kept the whole system charge neutral. The parameterization of ions implicitly included the first hydration shell around ions. The hydrated Na^+^ and Cl^-^ ions were given the “Qd” bead type with integral +1 and −1 charge, respectively. Details of the systems are listed in **Table S1**.

### Re-parameterization of PI(4,5)P_2_ Martini force field

The predominant form of PIP_2_ in the plasma membrane is PI(4,5)P_2_ with −4 charge, which indicates one of the phosphate groups being protonated (*43*). The current PIP_2_ model in Martini force field (residue name POP2) is based on PI(3,4)P_2_ with −5 charge. Hence, the charge on bead name P2 in Martini lipid POP2 is reduced from −2 to −1. Benchmark was done to compare the Martini CG PI(4,5)P_2_ model with the all-atom PI(4,5)P_2_ structure (residue name SAPI24) in the CHARMM36 lipid force field (*44*). Since CHARMM SAPI24 has one more double bond than the Martini POP2 model, the bead name C3A is modified to D3A with its type changing from C3 to C4. PyCGTOOL was used to check the correct CG to AA mapping (*45*). The new Martini PI(4,5)P_2_ model (POP5) is provided in **Table S2.** Bond length, angle pairs, and radius of gyration are calculated and compared for both the CG model and the atomistic model (**Table S3)**.

### CG simulation protocol and reverse mapping scheme

CG simulations in the current study are designed to simply allow faster convergence of membrane topology while keeping the secondary and tertiary structure of Piezo1 intact. Hence, the protein backbones were kept rigid using positional restraint with a force constant of 1000 kJ mol^-1^nm^-2^ and an elastic network (*46*) with a cutoff of 9 Å and a force constant of 500 kJmol^-1^nm^-2^. The convergence of the PIP_2_ lateral diffusion was monitored by the time evolution of lateral density maps of PIP_2_. **Figure S5** shows the PIP_2_s quickly diffuse towards Piezo1 within 1 μs and remain at the annular region of the protein throughout the 12 μs trajectory.

All the CG simulations were executed in GROMACS (version 2016.4) simulation package with the standard Martini v2.2 simulation setting (*47*). The protein and membrane systems were built using a modified enhanced version of the INSANE (INSert membrane) CG building tool. All lipid models and parameters used in this study follow the MARTINI v2.0 lipids, with the addition of the modified Martini PI(4,5)P_2_ model (POP5). The overall workflow of the simulations includes the initial construction of the Piezo 1 embedded membrane, energy minimization, isothermal-isochoric (NVT) and isothermal-isobaric (NPT) equilibration runs, and NPT production runs. Briefly, each system was firstly energy minimized (steepest descent, 5000 steps) without constraints. NVT simulations were carried out for 0.5 ns at 310.15 K with a timestep of 10 fs. A time step of 20 fs was used for the following NPT simulations. A cut-off of 1.1 nm was used for calculating both the electrostatic and van der Waals interaction terms; the potential-shift-Verlet algorithm was applied to take care of both interactions by smoothly shifting beyond the cutoff. Coulomb interactions were calculated using the reaction-field algorithm implemented in GROMACS. The neighbor list was updated every 20 steps using a neighbor list cutoff equal to 1.1 nm for short-range van der Waals. The temperature for each group (protein, membrane, ion, and water) was kept constant using the velocity rescale coupling algorithm with 1 ps time constant. For the NPT equilibration step, semi-isotropic pressure coupling was applied using the Berendsen algorithm, with a pressure of 1 bar independently in the cross-section of the membrane and perpendicular to the membrane with the compressibility of 3.0 × 10^−4^ bar^-1^. The pressure in newly built systems was relaxed in a 30 ns simulation using the Berendsen barostat with a relaxation time constant equal to 5.0 ps. Three-dimensional periodic boundary conditions were used. The production step for each system ran for 12 µs using Parrinello-Rahman barostat with a relaxation time constant of 12.0 ps. At the end of each CG simulation, the protein and lipids were mapped into its atomistic representation using Martini backward mapping scheme. The reverse-mapped atomic structures were solvated with CHARMM TIP3P water and 150 mM KCl using the CHARMM36 force field (*48*).

### AA simulation protocol

At the end of 12 μs CG-MD simulations, the reverse-mapped System I and II were truncated to 20.9×21.9×15.6 nm^3^ (System III) and 21.9×22.7×15.6 nm^3^ (System IV) in *xyz* dimensions, to reduce the all-atom system size to around 800,000 atoms. Two more systems (V and VI) were generated from the snapshot of System II (CG PC:PC/PIP_2_) at 12 μs in order to probe the role of PIP_2_ in Piezo1 gating. System V (PIP_2_-neutral) was prepared by removing all 39 PIP_2_ head group charges in GROMACS topology file from System IV (PC:PC/PIP_2_) and removed 156 K^+^ ions to neutralize the system. In system VI, instead of neutralizing the PI(4,5)P_2_ charge, all 39 PI(4,5)P_2_ were replaced by POPC. The all-atom systems were first minimized using 50000 steepest descent cycles in GROMACS (version 2016.4) package, and then underwent six stages of equilibrium run at 310.15 K using AMBER18 CUDA package as described in our previous Piezo1 simulation (*11*).

After equilibrium run on AMBER18, the systems were run on ANTON2 supercomputer with 2.0 femtosecond (fs) timestep. Lennard-Jones interactions were truncated at 11-13 Å and long-range electrostatics were evaluated using the k-Gaussian Split Ewald method (*49*). Pressure regulation was accomplished via the Martyna-Tobias-Klein (MTK) barostat, to maintain 1 bar of pressure, with a tau (piston time constant) parameter of .0416667 ps and reference temperature of 310.15 K. The barostat period was set to the default value of 480 ps per timestep. Temperature control was accomplished via the Nosé-Hoover thermostat with the same tau parameter. The *mts* parameter was set to 4 timesteps for the barostat control and 1 timestep for the temperature control. The thermostat interval was set to the default value of 24 ps per timestep. A half flat-bottom harmonic restraint with spring constant of 0.12 kcal mol^-1^ Å^-1^ was added between center of mass of the beams (residue 1339 - 1365) and the bottom of the pore (residue 2491-2546) to prevent the C-terminal of the beams drifting more than 30 Å away from the bottom of the pore in the absence of a loop sequence from the cryo-EM data. The amphiphilic helices (residue 1493 to 1553) were subjected to RMSD restraints for additional 90 ns at the beginning of Anton2 simulation to ensure the helical structures remain stable for the rest of microseconds production run. Due to Anton2 cluster processing capacity, all the systems at the end of 190 ns had to be cut into a smaller size (∼710,000 atoms) for longer production runs. Detailed information for the number of each component is indicated in **Table S4**.

### Ionic conductance simulations of WT and mutant Piezo1

In order to measure the ionic conductance, 16 POPC lipids from the outer membrane and 19 POPC lipids from the inner membrane were removed from the pore region. A total of 44 POPC lipids were removed to maintain the same ratio of leaflet surface. After removing the lipids in the pore, several 50 ns equilibrium simulations were conducted with and without membrane tension. It was found that a membrane tension of 14.2mN/m (around 10 bar) was necessary to remain the pore helices intact. Zero tension or higher membrane tension causes partial unfolding of the inner helices (**Figure S4**). Constant electric fields corresponding to the transmembrane potential of −250, −500, and −750 mV were applied perpendicular to the membrane to all the atoms in the simulation box, in the presence of symmetrical 150 mM KCl concentration and 14.2 mN/m membrane tension. In order to further validate the open state of Piezo1, the charge on three E2133 residues and three K^+^ ions were removed. The system was equilibrated for 20ns and the conductance was measured from three consecutive 50 ns simulations under a voltage of −500mV and a membrane tension of 14.2mN/m.

### PIP_2_ binding site analysis

The PIP_2_ binding and unbinding events are counted by GROMACS function ‘gmx select’, which print out whether the atom types PC PL of PIP_2_ headgroups are within the cut-off distance 5.7Å of carbon atom ‘name CZ or CE’ connecting with the charged groups of arginine/lysine residues of Piezo1 protein in 2μs AA simulation trajectory. The cut-off distance is the first minimum distance of the radial distribution function curve calculated between atom types PC PL of PIP_2_ headgroups and carbon atom name CZ CE of the arginine/lysine residues. Similar calculation is conducted for 12μs CG simulation trajectory, which prints out whether the bead name PO4 P1 P2 of PIP_2_ headgroups are within the cut-off distance 6.5 Å of bead name SC1 of arginine/lysine residues connecting to the charged groups. The percent of occupancy is calculated as the total occupancy time divided by the simulation time per each trajectory for each cationic residue.

### Whole-cell patch clamp electrophysiology

HEK293 cells were obtained from the American Type Culture Collection (ATCC) (catalogue number CRL-1573, RRID:CVCL_0045) and were cultured in minimal essential medium (MEM) (Life Technologies) containing 10% (v/v) Hyclone characterized fetal bovine serum (FBS) (Thermo Scientific), and penicillin (100 IU/ml) and streptomycin (100 µg/ml; Life Technologies). Cells were used up to 25-30 passages, when a new batch with low passage number was thawed. All cultured cells were kept in humidity-controlled tissue-culture incubator with 5% CO_2_ at 37°C. Cells were transiently transfected with cDNA encoding the mouse Piezo1 channel or its mutants tagged with GFP on its N-terminus in the pCDNA3 vector using the Effectene reagent (QIAGEN). Cells were then trypsinized and re-plated on poly-D-lysine-coated round coverslips 24 hours after transfection. Whole-cell patch clamp recordings were performed 36-72 hours after transfection at room temperature (22° to 24°C) as described previously (*14*). Briefly, patch pipettes were prepared from borosilicate glass capillaries (Sutter Instrument) using a P-97 pipette puller (Sutter instrument) and had a resistance of 4-7 MΩ. After forming gigaohm-resistance seals, the whole cell configuration was established, and the MA currents were measured at a holding voltage of −60 mV using an Axopatch 200B amplifier (Molecular Devices) and pClamp 10. Currents were filtered at 2 kHz using low-pass Bessel filter of the amplifier and digitized using a Digidata 1440 unit (Molecular Devices). All measurements were performed with extracellular (EC) solution containing 137 mM NaCl, 5 mM KCl, 1 mM MgCl_2_, 2 mM CaCl_2_, 10 mM HEPES and 10 mM glucose (pH adjusted to 7.4 with NaOH). The patch pipette solution contained 140 mM K^+^ gluconate, 1 mM MgCl_2_, 0.25 mM GTP, 5 mM EGTA and 10 mM HEPES (pH adjusted to 7.2 with KOH). Mechanical stimulation was performed using a heat-polished glass pipette (tip diameter, about 3 µm), controlled by a piezo-electric crystal drive (Physik Instrumente) positioned at 60° to the surface of the cover glass as previously described (*14*). The probe was positioned so that 10-µm movement did not visibly contact the cell but an 11.5-µm stimulus produced an observable membrane deflection. We applied an increasing series of mechanical steps from 12 µm in 0.5-µm increments every 5 s for a stimulus duration of 200 ms. The inactivation kinetics from MA currents were measured by fitting the MA current with an exponential decay function in pClamp, which measured the inactivation time constant (Tau). To calculate this time constant, we used the current evoked by the third stimulation after the threshold in the incrementally increasing step protocol in most experiments, except in cells where only the two largest stimuli evoked a current. In the latter case we used the current evoked by the largest stimulus, provided it reached 40 pA.

## ACKNOWLEDGMENTS

This work was supported by NIH Grants GM130834 (Y.L.L. and J.J.L.), NS101384 (J.J.L.), NS055159 (T.R.), GM093290 (T.R.), GM131048 (T.R.), F31-NS100484 (J.D.R.), F99-NS113422 (J.D.R.) and a WesternU intramural research award (Y.L.L. and J.J.L.). Computational resources were provided via the Extreme Science and Engineering Discovery Environment (XSEDE) allocation TG-MCB160119 (Y.L.L. and J.J.L.) and the Pittsburgh Supercomputing Center Anton2 allocations PSCA17006P-18007P (Y.L.L. and J.J.L.). The XSEDE program is supported by NSF grant number ACI-154862. The Anton2 machine at PSC was generously made available by D.E. Shaw Research and the Anton2 allocation program at PSC is supported by NIH Grant GM116961.

## AUTHOR CONTRIBUTIONS

Y.L.L, W.J., W.M.B-S., H.Z., and Y-C.L. designed and performed computer simulations; T.R., J.J.L., and J.S.D.R. designed and performed experiments; all authors analyzed data; Y.L.L., J.J.L., and T.R. designed the project and wrote the paper with inputs from all authors.

## SUPPLEMENTARY MATERIALS

**Video 1. A single potassium permeation event.** Trajectory of a permeating K^+^ ion during 9.5 ns simulation under −750 mV voltage. The backbone of the Piezo1 cap and pore domain is shown in orange. The DEED residues are shown in licorice with the atom color code (red oxygen, blue nitrogen, cyan carbon).

**Video 2. Accumulated potassium density along a multi-fenestrated pathway.** The isosurface of the K^+^ density calculated from 17 permeation events under −750 mV voltage. Density contours are shown at a level of 0.31 Å^-3^. The protein backbone is in cyan and hotspot residues in red licorice.

**Figure S1:**
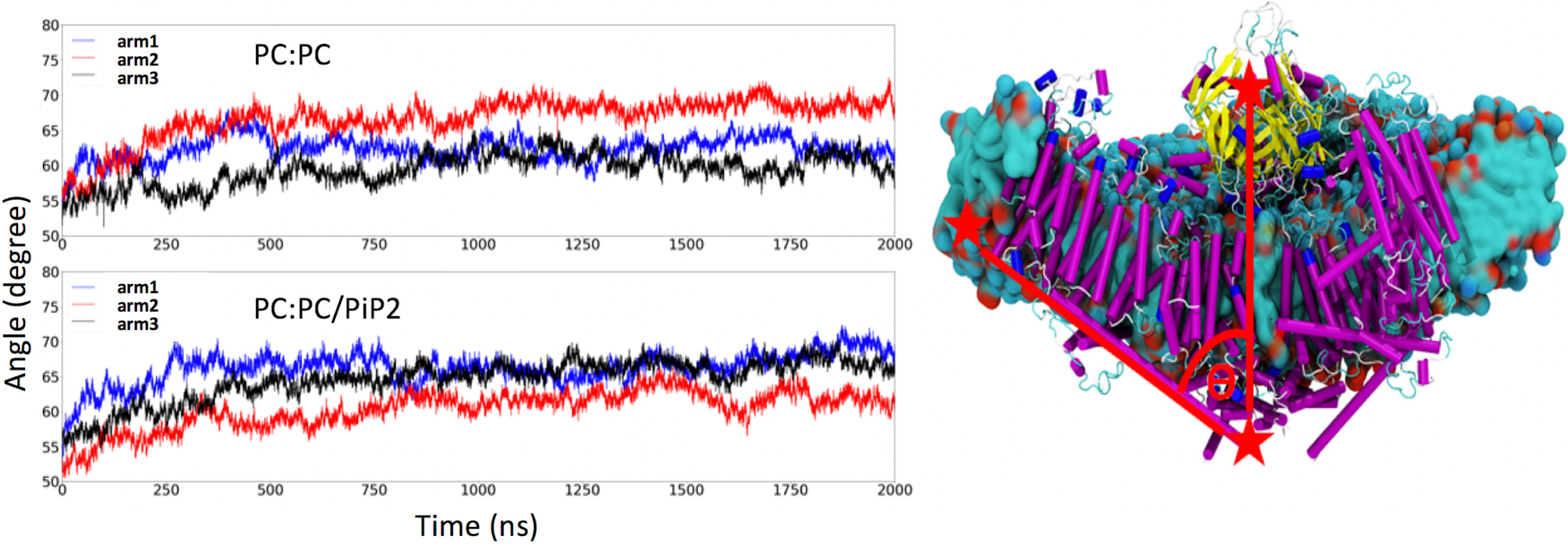
Angles of Piezo1 arms in PC:PC and PC:PC/PIP_2_ bilayer systems, showing the spontaneous flattening in crowded simulation environment.

**Figure S2:**
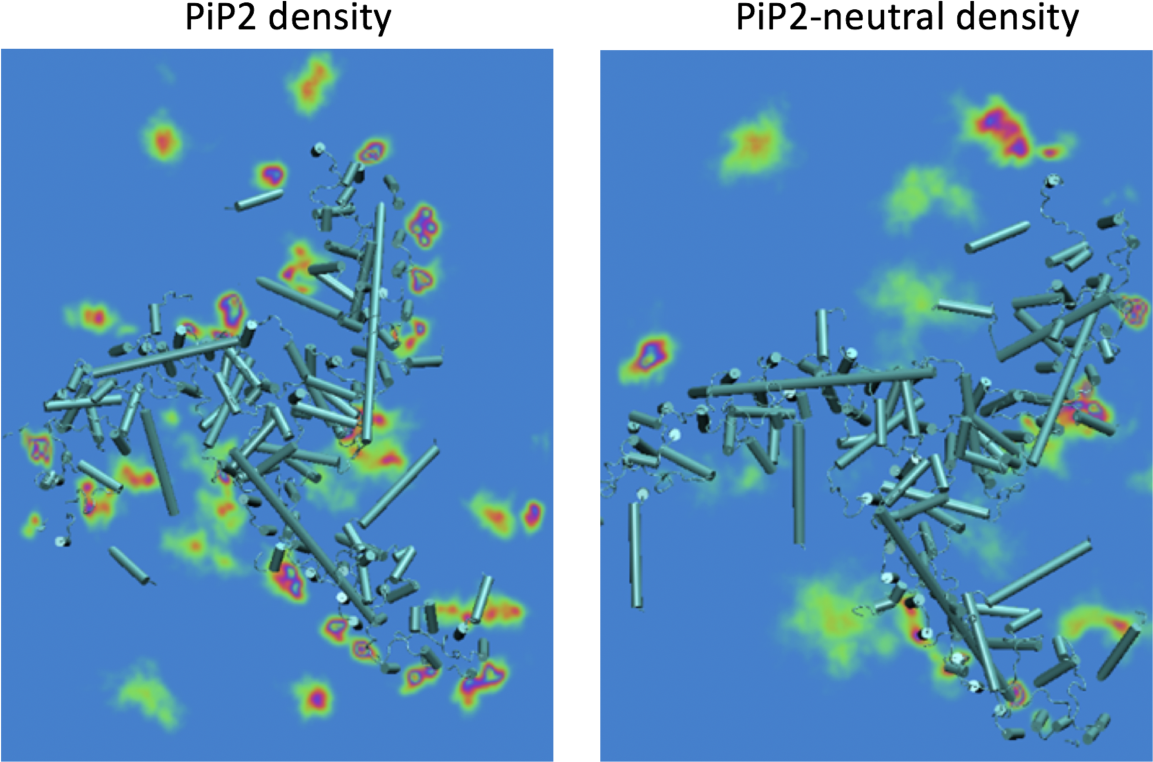
Average density of PIP_2_ lipids around Piezo1 (bottom view), calculated from the last 100 ns of 2 μS AA-MD simulations.

**Figure S3:**
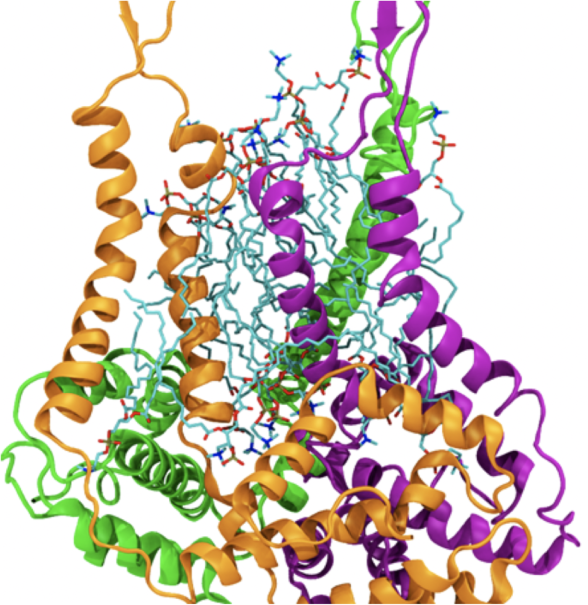
POPC lipids inside the Piezo1 pore in CG-MD simulations of Piezo1 in PC:PC/PIP_2_ bilayer. Lipids are shown in licorice with atom color code (red oxygen, blue nitrogen, cyan carbon, old phosphorus). The backbone of Piezo1 pore is shown in newcartoon mode with different colors for each subunit (orange, green, and purple).

**Figure S4:**
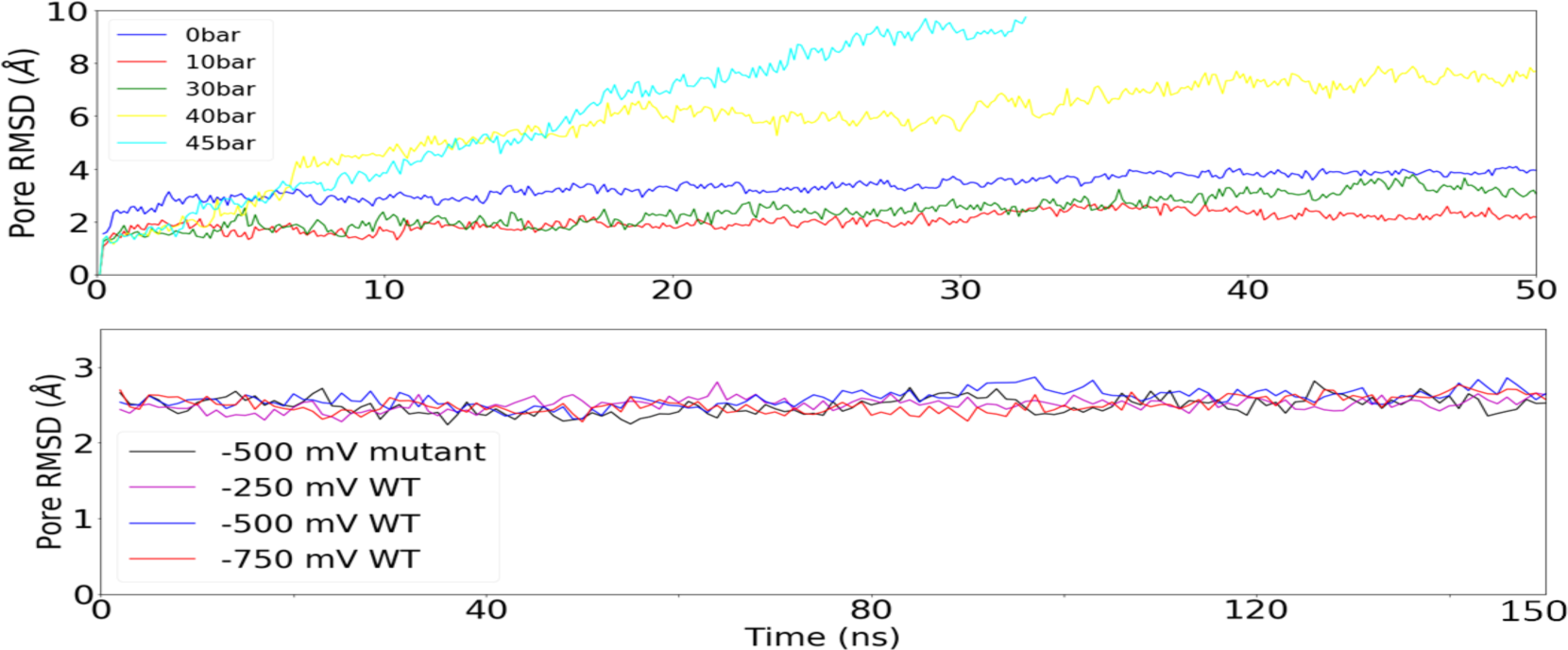
Root-mean-square deviation (RMSD) of inner pore helices of PIEOZ1 under different membrane tensions without voltage (top) and under different voltages with 10 bar tension (bottom).

**Figure S5:**
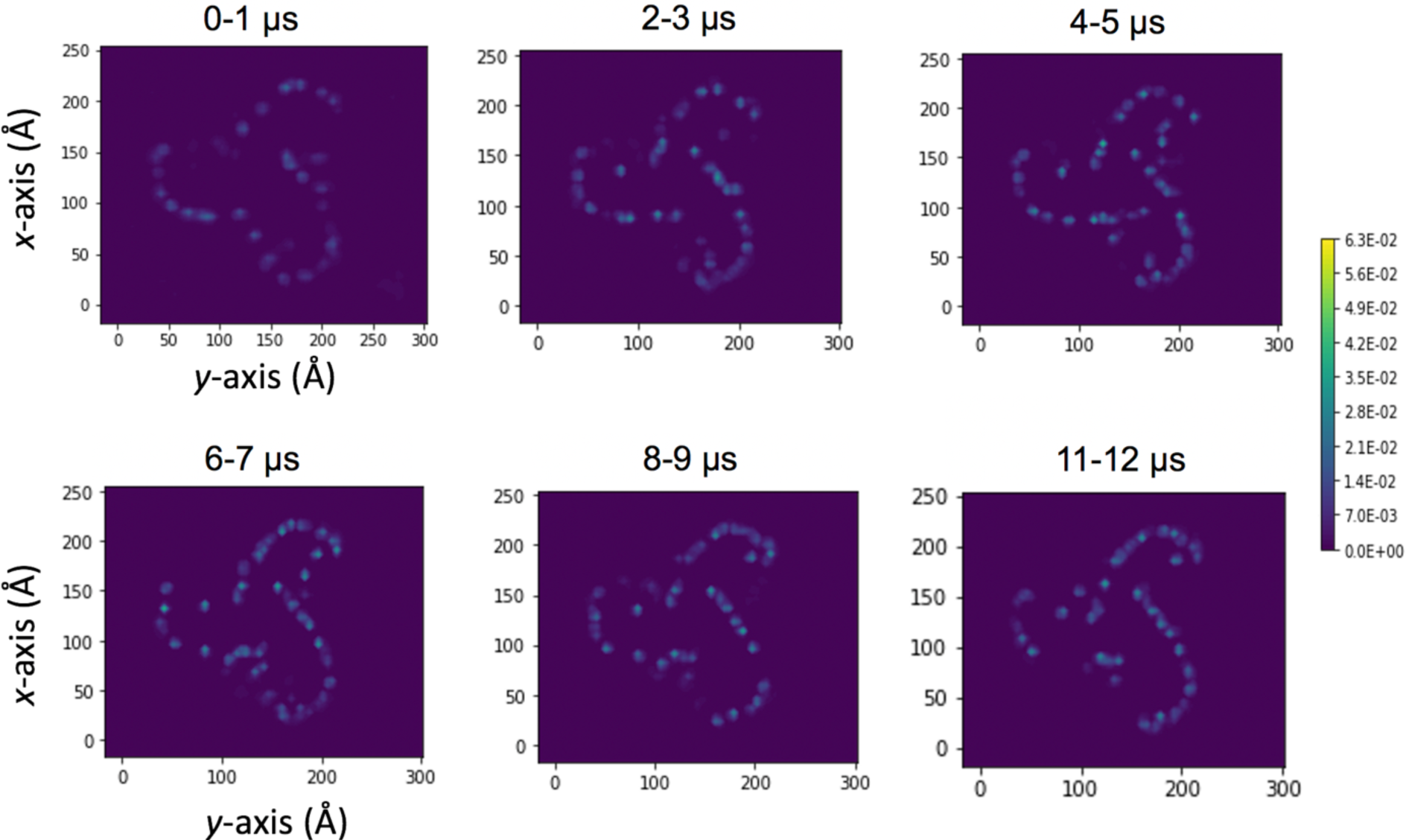
2D lateral density maps of PIP_2_ at lower leaflet over CG simulation trajectory.

**Table S1.** Details of CG Piezo 1 and AA Piezo 1 simulation systems.

| System |  |  | Inner |  | Outer | no. of |  |  |
| --- | --- | --- | --- | --- | --- | --- | --- | --- |
|  |  |  | PC | PIP2 | PC | Water | Na<sup>+</sup> | Cl<sup>+</sup> |
| CG | I | PC:PC | 872 | 0 | 952 | 109763 | 1174 | 1243 |
|  | II | PC:PC/PIP2 | 831 | 41 | 951 | 109414 | 1252 | 1157 |
| AA | III | PC:PC | 565 | 0 | 611 | 184522 | 590 | 759 |
|  | IV | PC:PC/PIP2 | 587 | 39 | 664 | 185688 | 799 | 712 |
|  | V | PC:PC/PIP2<br>neutral | 587 | 39 | 664 | 185688 | 643 | 712 |
|  | VI | PC:PC/PIP2<br>→PC | 626 | 0 | 664 | 186112 | 712 | 781 |

**Table S2a.** GROMACS topology file (bead type) for Martini PI(4,5)P2 model.

| nr | type | resnr | residue | atom | cgnr | charge |
| --- | --- | --- | --- | --- | --- | --- |
| 1 | P1 | 1 | POP5 | C1 | 1 | 0 |
| 2 | Na | 1 | POP5 | C2 | 2 | 0 |
| 3 | P4 | 1 | POP5 | C3 | 3 | 0 |
| 4 | Qa | 1 | POP5 | PO4 | 4 | -1.0 |
| 5 | Qa | 1 | POP5 | P1 | 5 | -2.0 |
| 6 | Qa | 1 | POP5 | P2 | 6 | -1.0 |
| 7 | Na | 1 | POP5 | GL1 | 7 | 0 |
| 8 | Na | 1 | POP5 | GL2 | 8 | 0 |
| 9 | C1 | 1 | POP5 | C1A | 9 | 0 |
| 10 | C4 | 1 | POP5 | D2A | 10 | 0 |
| 11 | C4 | 1 | POP5 | D3A | 11 | 0 |
| 12 | C1 | 1 | POP5 | C4A | 12 | 0 |
| 13 | C1 | 1 | POP5 | C1B | 13 | 0 |
| 14 | C1 | 1 | POP5 | C2B | 14 | 0 |
| 15 | C1 | 1 | POP5 | C3B | 15 | 0 |
| 16 | C1 | 1 | POP5 | C4B | 16 | 0 |

**Table S2b.**
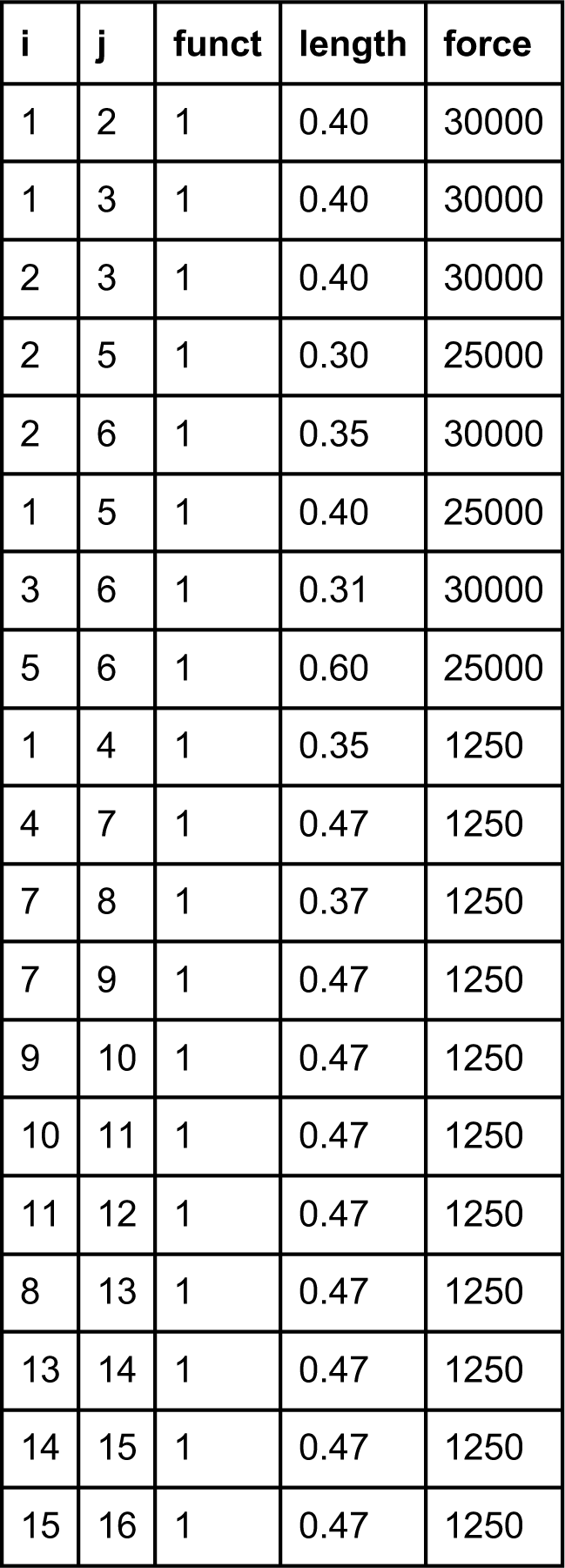
GROMACS topology file (bone length) for Martini PI(4,5)P2 model.

**Table S2c.** GROMACS topology file (angles) for Martini PI(4,5)P2 model.

| i | j | k | funct | angle | force |
| --- | --- | --- | --- | --- | --- |
| 1 | 4 | 7 | 2 | 140.0 | 25.0 |
| 7 | 9 | 10 | 2 | 180.0 | 25.0 |
| 9 | 10 | 11 | 2 | 100.0 | 10.0 |
| 10 | 11 | 12 | 2 | 120.0 | 45.0 |
| 8 | 13 | 14 | 2 | 180.0 | 25.0 |
| 13 | 14 | 15 | 2 | 180.0 | 25.0 |
| 14 | 15 | 16 | 2 | 180.0 | 25.0 |

**Table S2d.** GROMACS topology file (constraints) for Martini PI(4,5)P2 model.

| i | j | funct | length |
| --- | --- | --- | --- |
| 1 | 2 | 1 | 0.40 |
| 1 | 3 | 1 | 0.40 |
| 2 | 3 | 1 | 0.40 |
| 2 | 5 | 1 | 0.30 |
| 2 | 6 | 1 | 0.35 |
| 1 | 5 | 1 | 0.40 |
| 3 | 6 | 1 | 0.31 |

**Table S3a.**
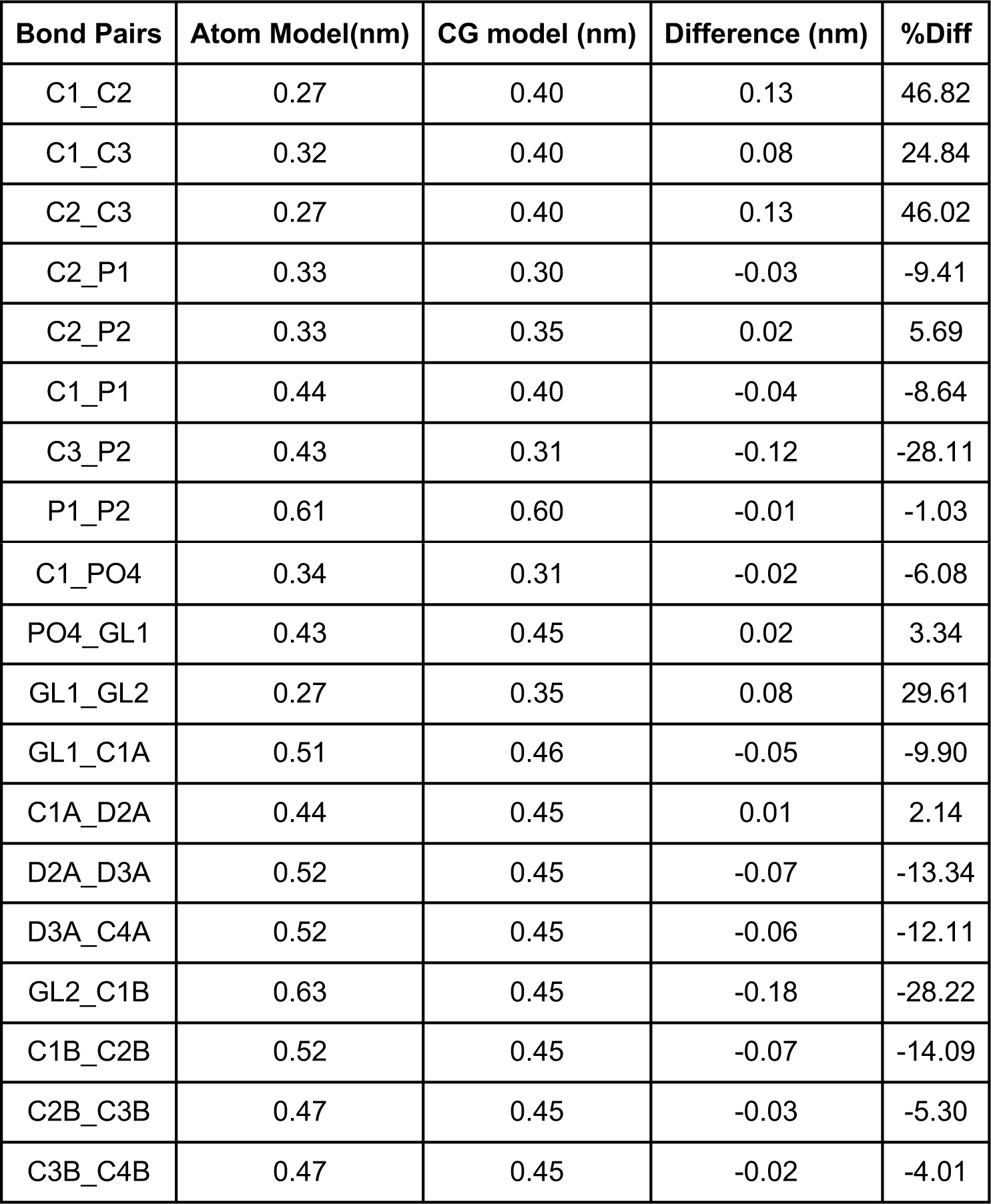
Bond length comparison between CG and AA model for PI(4,5)P2.

| Bond Pairs | Atom Model(nm) | CG model (nm) | Difference (nm) | %Diff |
| --- | --- | --- | --- | --- |
| C1_C2 | 0.27 | 0.40 | 0.13 | 46.82 |
| C1_C3 | 0.32 | 0.40 | 0.08 | 24.84 |
| C2_C3 | 0.27 | 0.40 | 0.13 | 46.02 |
| C2_P1 | 0.33 | 0.30 | -0.03 | -9.41 |
| C2_P2 | 0.33 | 0.35 | 0.02 | 5.69 |
| C1_P1 | 0.44 | 0.40 | -0.04 | -8.64 |
| C3_P2 | 0.43 | 0.31 | -0.12 | -28.11 |
| P1_P2 | 0.61 | 0.60 | -0.01 | -1.03 |
| C1_PO4 | 0.34 | 0.31 | -0.02 | -6.08 |
| PO4_GL1 | 0.43 | 0.45 | 0.02 | 3.34 |
| GL1_GL2 | 0.27 | 0.35 | 0.08 | 29.61 |
| GL1_C1A | 0.51 | 0.46 | -0.05 | -9.90 |
| C1A_D2A | 0.44 | 0.45 | 0.01 | 2.14 |
| D2A_D3A | 0.52 | 0.45 | -0.07 | -13.34 |
| D3A_C4A | 0.52 | 0.45 | -0.06 | -12.11 |
| GL2_C1B | 0.63 | 0.45 | -0.18 | -28.22 |
| C1B_C2B | 0.52 | 0.45 | -0.07 | -14.09 |
| C2B_C3B | 0.47 | 0.45 | -0.03 | -5.30 |
| C3B_C4B | 0.47 | 0.45 | -0.02 | -4.01 |

**Table S3b.** Angle comparison between CG and AA model for PI(4,5)P2.

| Angle Pairs | Atom Model (degrees) | CG model (degrees) | Difference (degrees) | %Diff |
| --- | --- | --- | --- | --- |
| C1_PO4_GL1 | 101.35 | 132.26 | 30.92 | 30.51 |
| GL1_C1A_D2A | 107.09 | 141.37 | 34.28 | 32.01 |
| C1A_D2A_D3A | 109.43 | 92.07 | -17.37 | -15.87 |
| D2A_D3A_C4A | 105.27 | 118.42 | 13.15 | 12.49 |
| GL2_C1B_C2B | 110.94 | 138.13 | 27.19 | 24.51 |
| C1B_C2B_C3B | 138.20 | 136.60 | -1.61 | -1.16 |
| C2B_C3B_C4B | 134.62 | 137.64 | 3.02 | 2.24 |

**Table S3c.** Radius of gyration comparison between CG and AA model for PI(4,5)P2.

| PI(4,5)P2 | Atom model | CG model | Difference | %Diff |
| --- | --- | --- | --- | --- |
| Radius of Gyration(nm) | 0.77 | 0.81 | 0.04 | 5.28 |

**Table S4.** Details of extended all-atom Piezo 1 simulation systems.

| System | Inner |  | Outer | no. of |  |  | Time (ns) |
| --- | --- | --- | --- | --- | --- | --- | --- |
|  | POPC | PIP2 | POPC | Water | K<sup>+</sup> | Cl<sup>+</sup> |  |
| III-ext | 545 | 0 | 602 | 155033 | 623 | 692 | 2000 |
| IV-ext | 537 | 38 | 607 | 160133 | 695 | 612 | 2000 |
| V-ext | 517 | 37 | 593 | 151401 | 605 | 674 | 1810 |
| VI-ext | 508 | 0 | 566 | 144154 | 583 | 652 | 810 |

